# VIRUS-MVP: A framework for comprehensive surveillance of viral mutations and their functional impacts

**DOI:** 10.1101/2025.02.16.636315

**Authors:** Muhammad Zohaib Anwar, Ivan S. Gill, Madeline Iseminger, Anoosha Sehar, Emma J. Griffiths, Damion Dooley, Jun Duan, Khushi Vora, Gary Van Domselaar, Fiona S.L. Brinkman, Paul M. K. Gordon, William W.L. Hsiao

## Abstract

As viruses evolve, they accumulate genetic mutations that can influence disease severity, transmissibility, and the effectiveness of vaccines and therapeutics. Real-time tracking of viral mutations and their functional impacts is essential to understand these changes and assess their implications for public health responses. VIRUS-MVP is an interactive, portable platform designed for the comprehensive surveillance of viral mutations. Initially developed for SARS-CoV-2, it now fully supports mpox and is expanding to include influenza and RSV. The platform links viral mutations to functional annotations, providing insights into their predicted effects on viral infectivity, immune evasion, and protein functionality. It features an interactive interface for visualizing mutation distributions, a modular and reproducible genomics workflow, and a curated annotation resource that captures known impacts on viral proteins and host interactions. Users can also import custom functional annotations to tailor analyses to specific research needs or emerging pathogens. Developed collaboratively with public health and academic partners, VIRUS-MVP enhances understanding of viral evolution and its public health impact by bridging genomic data with biological insights. The platform is open-source, adaptable, and accessible on GitHub.

## 1. Introduction

During the COVID-19 pandemic, genomic epidemiology offered unprecedented insights into the evolution and transmission dynamics of SARS-CoV-2, the virus responsible for the pandemic. In previous viral outbreaks, genomic epidemiology was mainly applied retrospectively, i.e., employing genomic data for analyses conducted after the events, or in real-time for a specific region (1–3). However, the advent of COVID-19 marked a paradigm shift, enabling real-time surveillance due to the unprecedented scale of genomic data generation, sharing, and analysis (4, 5). This transition underscores the transformative potential of genomic epidemiology in responding to pandemics. As the world navigates the post-pandemic landscape, the importance of continued surveillance for circulating viruses like influenza, mpox, and respiratory syncytial virus (RSV), alongside monitoring for emerging pathogens, cannot be overstated (6, 7). The global increase in public health genomics capacity achieved during the COVID-19 pandemic positions us to confront future infectious disease threats more effectively. As part of that capacity building, we developed COVID-MVP (8), a platform designed to monitor SARS-CoV-2 population structures, mutations, and real-time functional impacts of mutations. COVID-MVP bridged a significant gap in genomic surveillance and public health response by connecting the mutations harboured by variants to their possible functional impacts (8).

RNA viruses exhibit moderately high mutation rates due to the error-prone nature of their replication machinery (9). The accumulation of mutations leads to a diverse viral population, allowing the viruses to adapt to changing environments and potentially evade host immune responses. During the COVID-19 pandemic, several visualization dashboards and analytical workflows were developed to track the prevalence of mutations; popular examples include CovMT (10), CoV-Spectrum (11), CovRadar (12), and Outbreak.info (13) (10–13). Such tools provide valuable insights into the mutational landscape of SARS-CoV-2, summarizing mutations by geographical location, sampling date, and disease severity. They also offer data on variant proportions, common mutations, and associated hospitalization and mortality probabilities. COVID-MVP distinguished itself from other mutation-tracking tools by connecting mutations to their functional impacts in real time. This capability was crucial for assessing the evolving viral population and epidemiological characteristics, such as transmissibility, disease severity, immune escape, and the effectiveness of vaccines and therapeutics (14).

The need for ongoing genomic surveillance becomes increasingly paramount as we transition from pandemic response to preparedness. Building upon the development of COVID-MVP, we here present VIRUS-MVP, a virus-agnostic platform comprising three standalone components, each contributing to the platform’s flexibility, portability, and adaptability. The visualization component is a user-friendly interactive webpage powered by Python (using Plotly and Dash) and JavaScript. The second component is the genomics workflow, implemented in the Nextflow workflow management framework (15) and following the nf-core guidelines (16). The workflow processes genomic data in a modular, reproducible, and scalable manner, offering a plug-and-play approach for customization and selection of modules based on pathogen characteristics. Finally, the third component is the functional annotation database, named Pokay. Pokay is a manually curated database integrated into the genomic variant data to annotate the mutations and their functional impact.

Given the high mutation rates in many viruses and the need to identify rare alleles, a tool like VIRUS-MVP is essential for ongoing genomic surveillance and public health response. VIRUS-MVP can be tailored for different viruses, catering to each pathogen’s specific characteristics. This design enables real-time monitoring of population structures, mutations, and their functional impacts. By harnessing the power of genomic epidemiology, VIRUS-MVP represents a significant advancement in our ability to understand viral sequencing data and helps public health practitioners, epidemiologists, and academic researchers by quickly summarizing the functional significance of viral mutations. All three components of the framework are available open-source (see *Availability and Deployment*) under the Massachusetts Institute of Technology (MIT) and GPL-3.0 licenses.

## 2. Materials and Methods

### 2.1 Genomics Data and Formats

The VIRUS-MVP genomics workflow accepts assembled contigs (e.g., from clinical specimens) or paired-end reads (e.g., from wastewater sampling) along with an optional metadata file (in CSV format) and outputs a collection of identified mutations as a Genome Variant Format (GVF) file. The metadata file can include multiple variables such as sampling date, geographic location, and/or lineage information. These variables can be used as criteria for grouping samples. Currently, the genomics workflow supports processing genomic data from two viruses: SARS-CoV-2 and mpox.

The workflow supports the ability to choose from various compatible tools for a given step; for instance, more than one variant calling tool is available to choose from based on the sequence data type and sequencing depth (17). This flexibility, however, adds interoperability challenges as different tools have different output formats. To address this challenge, we use standardized data formats and have developed helper functions to convert the varying analytic tool output formats into these standard formats. We have used VCF (Variant Call Format) and GVF (Genome Variation Format) files as the standard formats for variant data. VCF is a widely adopted community standard for representing genomic variants, thus promoting data interoperability and exchange. GVF provides a structured encoding of genomic variations along with associated functional annotations.

We are developing VIRUS-MVP to support additional specific pathogens such as RSV and influenza. Additionally, VIRUS-MVP’s modular architecture and developer documentation allow end users/developers with modest coding expertise to adapt the workflow to other viruses rapidly.

### 2.2 Annotation Sources

VIRUS-MVP can utilize multiple data sources to annotate viral mutations. Primary data sources include

#### NCBI

The genomics workflow utilizes NCBI GenBank (18) to annotate each virus’s genomic features (including genes and coding sequences (CDS), non-coding regions, and protein products), adding context to the mutations they harbour.

#### Problematic Sites Identification

Sequencing errors, contamination, and hypermutable sites must be identified to ensure the accuracy of downstream analyses, such as phylogenetic reconstruction, variant interpretation, and genomic epidemiology. We have incorporated information regarding problematic sites identified in SARS-CoV-2 genomes using data from De Maio *et al.* (*19*). These observations are marked as either ‘Mask’ or ‘Caution’ based on the severity of their impact on the analysis. Each site is further labeled with a more specific potential problem. The problematic sites are recorded and available via our GitHub repository (20). A custom Python script is integrated into the workflow to parse the observations and integrate them with the mutations observed in the sequencing data analyzed.

#### Pokay Functional Database

At the heart of this framework is the continuous curation of functional annotations and their connection to mutations. This process is essential for understanding how genetic changes impact the behaviour of viruses, aiding in our comprehension of their evolution and disease-causing mechanisms. For the COVID-19 pandemic response, one of our primary goals was to curate functional annotations for mutations in the SARS-CoV-2 genome. Since then, Pokay has also been extended to mpox and influenza annotations, broadening its scope to encompass a wider range of viral pathogens.

For COVID-19 surveillance, Pokay has served as one of the most comprehensive resources for understanding the functional implications of SARS-CoV-2 mutations. Pokay is hierarchically structured, with functional categories paired with specific features of the viral genome. For instance, categories such as “S_convalescent plasma escape” describe functions related to convalescent plasma escape specifically associated with mutations in the SARS-CoV-2 S gene (encoding the spike protein). Similarly, categories like “RdRp_pharmaceutical_effectiveness.txt” capture mutations in the RdRp mature peptide associated with pharmaceutical effectiveness. Within each functional category, specific functions and associated mutations are curated. Additionally, comprehensive information, including the source(s), citation(s), and URL(s) of relevant studies, is listed to facilitate further exploration and verification.

Pokay’s hierarchical design disentangles the complex relationships between mutations and their functional consequences. Notably, a mutation may be implicated in single or multiple functions, and conversely, a single functional impact may be associated with individual mutations or combinations thereof. A Python script has been provided with the repository to merge functional data from all categories within Pokay, generating a structured TSV file, which is then used for integrating functional information into the GVF files, facilitating visualization and surveillance reporting. Pokay comprises over 40 unique functional categories and contains more than 4000 functional annotations for over 1000 mutations, individually or in combination. Pokay is updated regularly with the latest information on mutations and their associated functions. This undertaking involves a comprehensive literature search and curation to ensure that Pokay remains a reliable and up-to-date resource for researchers and stakeholders.

Furthermore, we are working on ontologizing the functional annotations, mutations, and additional data using a controlled vocabulary. This ongoing effort is the subject of a separate study, which will also include our collaboration with the Coronavirus Infectious Disease Ontology (CIDO) to integrate all mutations into CIDO (21, 22). This initiative aims to enhance interoperability, data integration, and semantic consistency across various resources.

### 2.3 Data Standards

We adhere to a range of established genomic data standards to ensure consistency, interoperability, and accuracy in data representation, analysis, and sharing, including, as a notable example, the use of Human Genome Variation Society (HGVS) nomenclature (23). The HGVS format provides a standardized and internationally recognized method for describing mutations, facilitating precise annotation and interpretation of mutations. The adoption of HGVS ensures that genomic variants are uniformly and consistently represented across datasets and analyses, facilitating cross-sector integration and interpretation (24).

A key guiding principle in VIRUS-MVP’s development is adherence to the FAIR (Findable, Accessible, Interoperable, and Reusable) data principles (25). By making genomic data and its associated metadata findable through standardized ontologies, accessible through open data portals, interoperable via structured formats like JSON and YAML, and reusable by ensuring transparent data provenance, VIRUS-MVP enables robust data sharing and reusability. These principles not only support scientific reproducibility but also facilitate collaboration across diverse research communities and public health agencies.

## 3. Results

### 3.1 Implementation

Here we describe the implementations of the components that comprise VIRUS-MVP and how they function independently and together as a system.

#### 3.1.1 Visualization Design

The visualization component is designed to be customizable as the visualization is rendered by the configuration file produced as part of the genomics workflow. Visualization is implemented almost entirely in Python (using the graphing library Plotly and the visualization framework Dash) and is supported by Javascript components. Currently, two different implementations are available: one supports real-time processing of user-uploaded data (see the Design of the Genomics Workflow below), enhancing its suitability for data analysis and dashboarding, while the other one without the upload function is more suitable for visualizing the pre-processed data for public-facing surveillance dashboards. Both versions are available for local and cloud deployment and can be cloned from the GitHub repository (see *Availability and Deployment*).

##### Heatmap

The visualization component is an interactive web application centered on a heatmap that encodes the frequency of mutations at a given site across different groups (e.g., SARS-CoV-2 lineages, as shown in Figure 1 or user-defined grouping). The heatmap has multiple axes, encoding the following attributes: **Top:** nucleotide position of mutation with respect to the reference viral genome e.g., NC_045512.2 for SARS-CoV-2, and NC_063383.1 for mpox (or user-provided reference genome).

**Figure 1.**
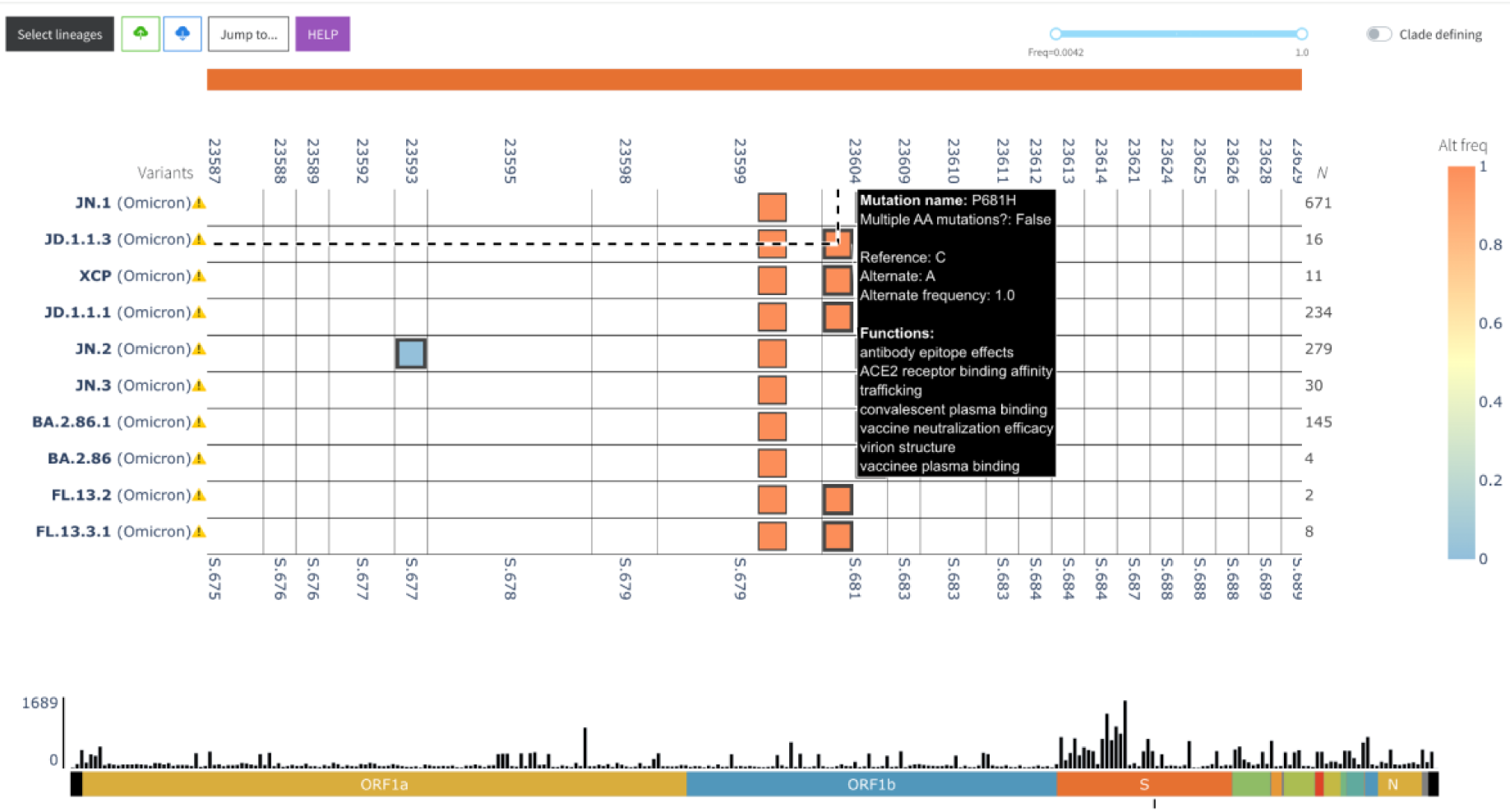
Screenshot of VIRUS-MVP web interface with SARS-CoV-2 sequences grouped into lineages. The central heatmap encodes mutation frequency with functional information available on hover. The bottom histogram shows the distribution of mutations across the genome.

**Bottom:** amino acid position of the mutation within its gene, or if the mutation is in an intergenic region, the number of amino acid positions upstream/downstream of the closest gene.

**Left:** sample ID (wastewater mode) or sample groups based on different criteria e.g., time, region, or lineage. In the case of lineages, variant status is also added to the group, e.g., Variants of Concern (VOC). **Right:** Number of sequences or paired-end reads in each group.

Heatmap cells encoding insertion and deletion mutations are annotated with unique markers, and cells representing mutations with known functions are represented with a thicker border. Users can hover over the heatmap cells for more information on each mutation. If the mutations have associated functions, users can also click on the cells to open a modal with a description and source of each function, as shown in Figure 2.

**Figure 2.**
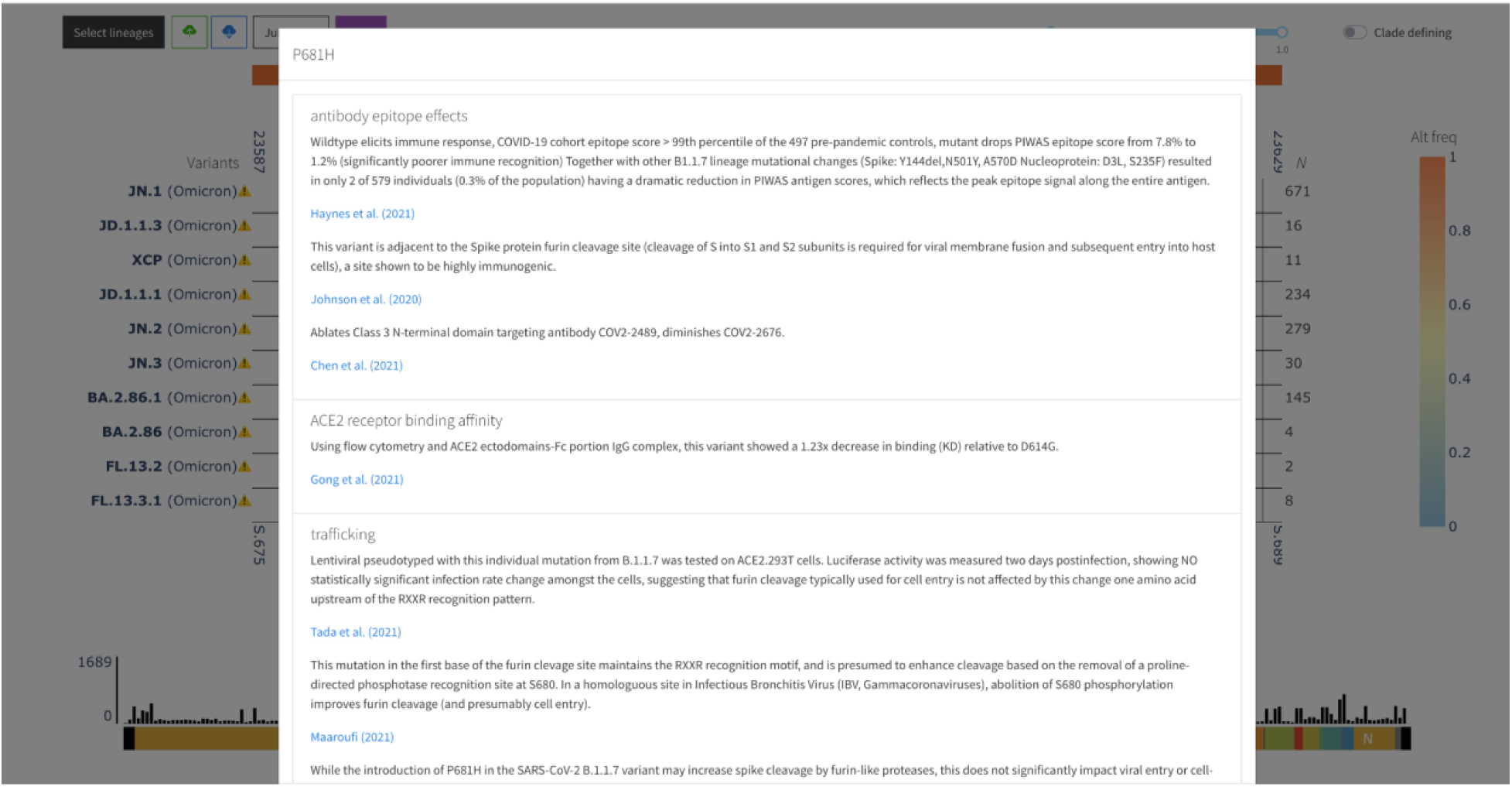
The modal opens when a user clicks on a heatmap cell with functional annotations in VIRUS-MVP. A description of each function and its primary source are provided as clickable links.

##### Histogram

Under the heatmap, a histogram encodes the total number of mutations across all visualized groups per 100 nucleotides with respect to the reference. This secondary visualization summarizes the distribution of mutations across the genome, as shown in Figure 1.

##### Editing the Visualization

The visualization can be edited in several ways. A “select groups” button at the top-left of the application opens a modal that allows users to filter or change the order of groups/lineages visualized in the heatmap. A “clade-defining” switch enables users to quickly filter out mutations with <75% frequency. We define the clade-defining mutations similar to the Outbreak.info (13). However, a frequency slider to the left of this switch provides greater granularity for filtering mutations by frequency if necessary.

##### Downloading and Uploading Data

An “upload” button to the right of the “select groups/lineages” button allows users to upload known variants as a VCF file. Alternatively, users can upload a collection of sequences as a FASTA file. The genomics workflow processes the input data and presents the result alongside the existing data as a heatmap (currently supported for SARS-CoV-2 only). A “download” button is also available that allows users to export a ZIP file containing the surveillance reports for each group (see *Surveillance Reports* section below).

#### 3.1.2 Functional Annotation Framework

In addition to the Pokay function database already described, we have developed a framework that allows the integration of annotations from additional sources. This process required the development of a standardized, generic template that allows external annotations to be integrated with variant calls, either in conjunction with or in place of Pokay annotations. The development of this template posed significant challenges, including the need for extensive data standardization, controlled vocabulary, and consistent formats, such as the Human Genome Variation Society (HGVS) nomenclature. Standardizing the format and vocabulary of external annotations was crucial to ensure interoperability and compatibility with existing genomic analysis pipelines. This process involved meticulous data curation and harmonization efforts to reconcile differences in annotation styles and terminology used across different sources.

To facilitate the integration of external annotations, we developed a Mutation Functional Annotation Contextual Data Specifications package (see *Availability and Deployment* section). Users can use this template to add functional information using a controlled vocabulary and pick lists and integrate them into their analyses. To validate user-developed annotations, we have developed a DataHarmonizer (26) template for VIRUS-MVP functional annotation. This harmonization process enhances data quality and creates interoperability between different systems, facilitating collaborative research efforts and cross-study comparisons.

#### 3.1.3 Design of the Genomics Workflow

The genomics workflow is implemented in the Nextflow workflow management system (15), following the nf-core guidelines (16). The workflow is developed in a modular structure where each process or step of the analysis is implemented as a discrete module. Each module is containerized using software container technologies, including Singularity (27, 28), Docker (29), and BioConda (30). The modules are connected in a particular order to form sub-workflows. The workflow contains more than 50 modules and 10 sub-workflows that perform a diverse set of analyses. The order and execution of these modules and sub-workflows are controlled by a main script tailored to each virus. The core parameters to run the workflow can be supplied in a YAML configuration file or provided directly on the command line. These parameters control the types of analyses to run and the arguments for a given analysis. At a minimum, the user needs to provide a FASTA file containing viral sequences or a VCF file as input. An accompanying metadata file containing information about the samples can be provided to assist in grouping samples based on user-defined criteria. A detailed structure of the workflow is presented in Figure 3.

**Figure 3.**
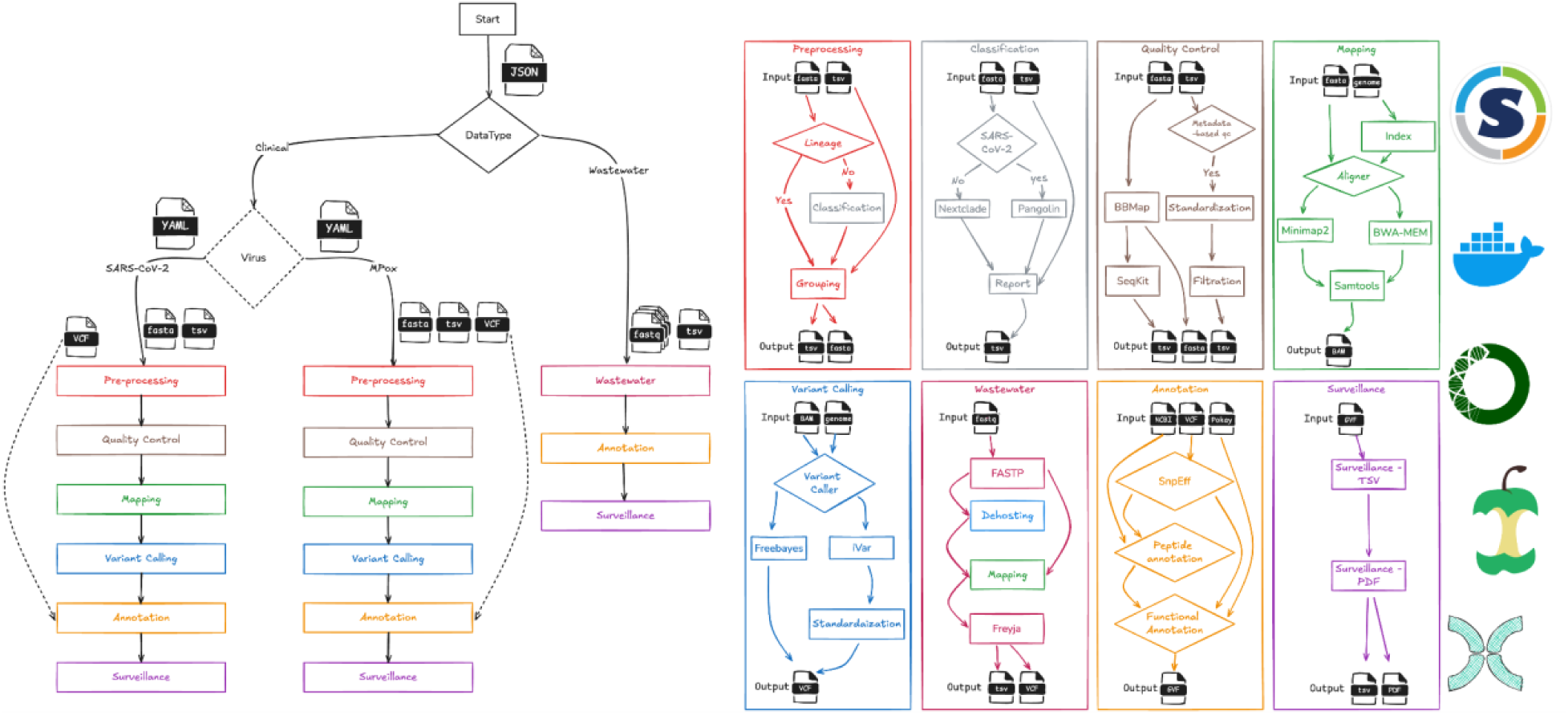
Design of genomics workflow. The right panel of the image shows a library of sub-workflows. Containerized environments and bioinformatics frameworks ensure modularity and reproducibility. The left panel displays how these sub-workflows are connected and controlled to process data from different viruses and data types, such as wastewater. The YAML and JSON files encode the parameters for sub-workflows.

Before variant-calling, the user can group samples by lineage/clades (default) or other metadata variables such as time, geographical location, or any other categorical variable in the supplied metadata file. If the lineage/clade information is not provided in the metadata file, the user can classify the samples using Pangolin (31) and Nextclade (32). Pangolin is specifically designed for SARS-CoV-2, whereas Nextclade can be used for different viruses. Both tools are integrated into the workflow, and an in-house Python script is used to merge the classification report with the metadata file for downstream analysis.

To filter out low-quality or missing data, the workflow allows the user to perform quality filtering based on the grouped samples with (i) metadata-based filtering and (ii) sequence-based filtering. Several Python scripts are embedded in the workflow to perform metadata-based filtering. We also provide recommended default cut-off filters derived from our experience developing COVID-MVP. The script allows options to filter samples that (i) originate from non-human hosts; (ii) have a length shorter than the user-specified length (default: 95% of the length of reference sequence); (iii) lack a complete sample collection date (i.e., YYYY-MM-DD); and (iv) were collected outside the user-defined start and end dates (default: no limits). For the sequence-based quality filtering, the workflow uses two open-source tools, BBMAP (33) and SEQkit (34), that can filter out low-quality sequences, especially based on missing information (number of N bases).

We perform the variant calling to characterize the virus’s population structure. This process includes mapping sequences against the default reference genome (or provided by the user). The mapping process is carried out by one of the two open-source tools available in the workflow, minimap2 (35) and BWA-MEM (36). The resulting BAM file is then used to call variants. Two variant calling tools are currently available in the workflow, Freebayes (37) and iVar (3). Freebayes produces a Genomic Variant Call Format (GVCF) file, which is further processed using an in-house Python script to filter out low-quality mutations and output a VCF file. For iVar, which produces a TSV file, we converted it to a VCF file using an in-house Python script. We have integrated a dedicated sub-workflow for processing the paired-end short reads to perform a similar analysis on the environmental samples (wastewater). The sub-workflow uses FastP (38) to trim/filter out low-quality reads. Like WGS sequences, quality-controlled reads are mapped to the reference genome using BWA-MEM or minimap2, resulting in a BAM file. Unlike WGS sequences, the wastewater samples may contain multiple co-circulating lineages/clades for a given virus. The workflow uses iVar to trim primers and call variants relative to the reference alignment file. This workflow embeds the Freyja workflow (39) to recover relative lineage abundances from mixed viral samples. Freyja is an open-source tool that uses lineage-determining mutational barcodes derived from the UShER global phylogenetic tree as a basis set to solve the de-mixing problem. It produces a TSV file which is then converted to a VCF file using an in-house Python script.

The workflow uses several annotation sources, including the Pokay functional database, to profile and summarize mutations present in the supplied viral sequence data. The annotation sub-workflow has several modules and processes, some of which are used universally, while others are specific to the virus under study. One of the universally used tools is SnpEff (40), which is used to summarize mutations.

SnpEff summarizes each mutation, including position, gene name, gene ID, and feature type. Variants and their annotations are recorded in GVF files. Multiple in-house Python scripts are embedded in the workflow to parse and annotate NCBI and RefSeq database records (41). The GVF file is also annotated using the Pokay functional database and/or any other annotations formatted using the template provided (see the *Functional Annotation Framework* section above). The resulting annotated GVF file is then used for visualization.

In addition to the GVF files produced by the workflow for visualization, the workflow also generates summarized surveillance reports. These surveillance reports are designed in two different formats: (i) A TSV file that contains each identified mutation and its corresponding functions and additional contextual information; and (ii) a PDF file containing human-readable summary tables. The TSV file can be used as input for integrating VIRUS-MVP with other surveillance and downstream analysis tools. Each column’s description, type, and source in the TSV file are available in our GitHub repository (see the *Availability and Deployment*). The surveillance reports are intended for end users who wish to incorporate them directly into their genomic surveillance reporting processes. The PDF file includes three sections ( Figure 4): the first section, *Information on the variant or cluster of sequences section* includes (i) the range of dates for genomes based on the metadata field “*sample collection dates*” that are either provided by the user or extracted from the dataset; (ii) the number of genomes used in the analysis, to give context for the frequencies provided in section 3, and (iii) a list of Pangolin lineages or Nextclade assigned clades identified in the dataset. The second section *(Indicator section)* contains a three-column table of disease indicators for public health surveillance, e.g., *transmissibility between humans, infection severity, etc.* (indicators can be specified for each run in an input file: see the *Availability and Deployment*), functional categories (from Pokay) grouped by disease indicators (e.g., *ACE2 receptor binding affinity, viral load*, and *outcome hazard ratio* are classified under *infection severity)* and the mutations identified for each category and indicator. Finally, the third section (*Mutation significance section)* consists of a table that provides each mutation’s prevalence, functional impact, hyperlinked citation for the study, lineage, number of sequences in the lineage from the dataset, reference allele, alternate allele, and alternate frequency. An example of a surveillance report in PDF format is available in the nf-ncov-voc repository (see the *Availability and Deployment*).

**Figure 4.**
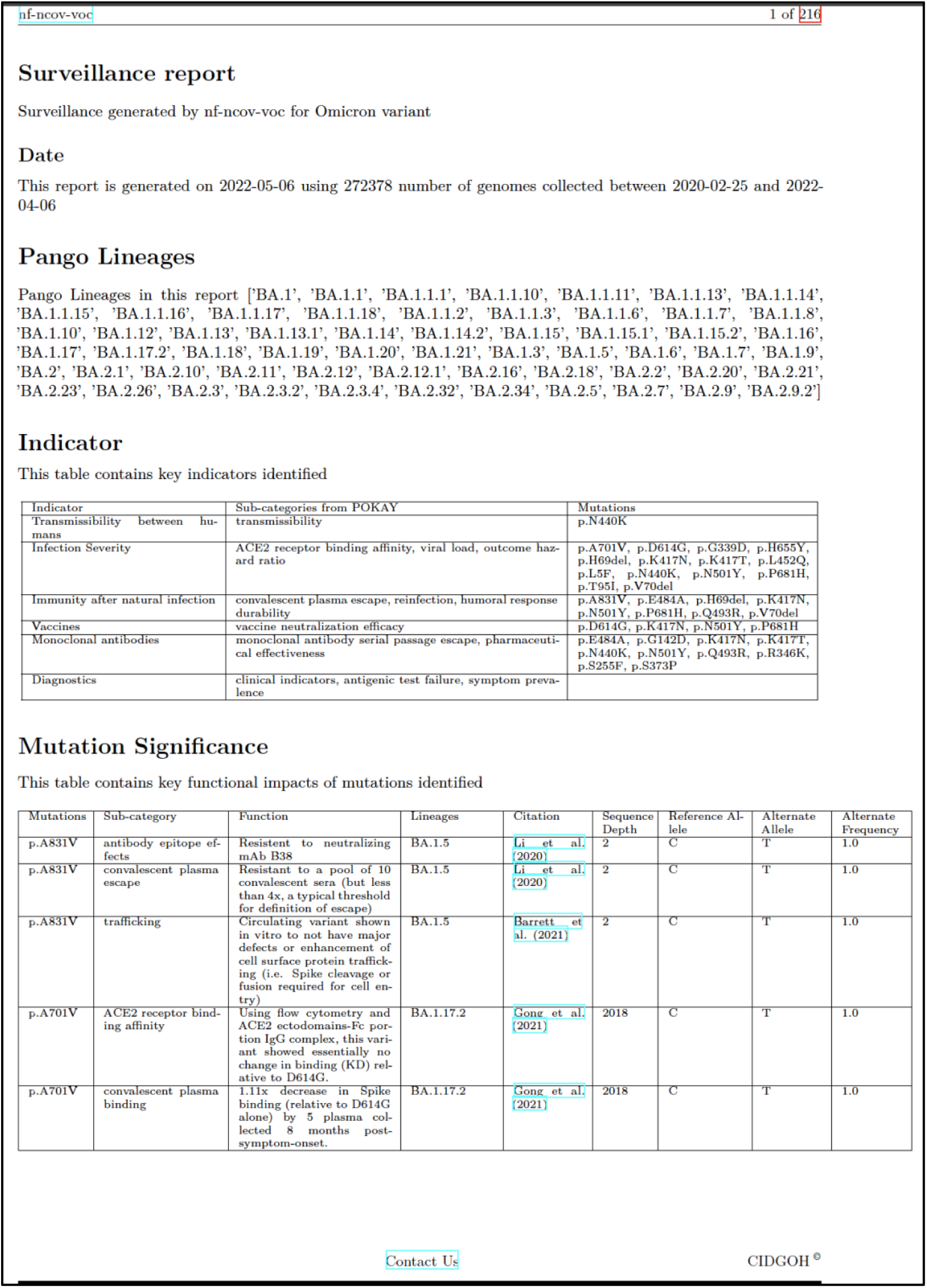
An example SARS-CoV-2 surveillance report with information on the number of genome sequences, identified viral (Pango) lineages, a table highlighting mutations associated with high-level surveillance indicators, and a table summarizing each mutation’s significance.

### 3.2 Use cases

VIRUS-MVP’s flexible workflows and customizable datasets allow it to answer different questions related to viral evolution and genomic epidemiology. Here, we demonstrate two applications of VIRUS-MVP by leveraging data from the SARS-CoV-2 dedicated instance, which uses the Canadian national SARS-CoV-2 genomic surveillance dataset.

#### 3.2.1 Longitudinal analysis

The custom grouping functionality of genomics workflow enabled us to perform a longitudinal analysis by organizing the dataset by epi-week rather than by lineage (Figure 5). This approach allowed us to perform a week-by-week assessment of mutation frequency across the sample population. With adaptable axis settings in the visualization component, we could dynamically relabel the Y-axis with epi-week on the left and the number of isolates per week on the right. This flexibility facilitated the monitoring of significant mutations over time, independent of lineage assignments.

**Figure 5.**
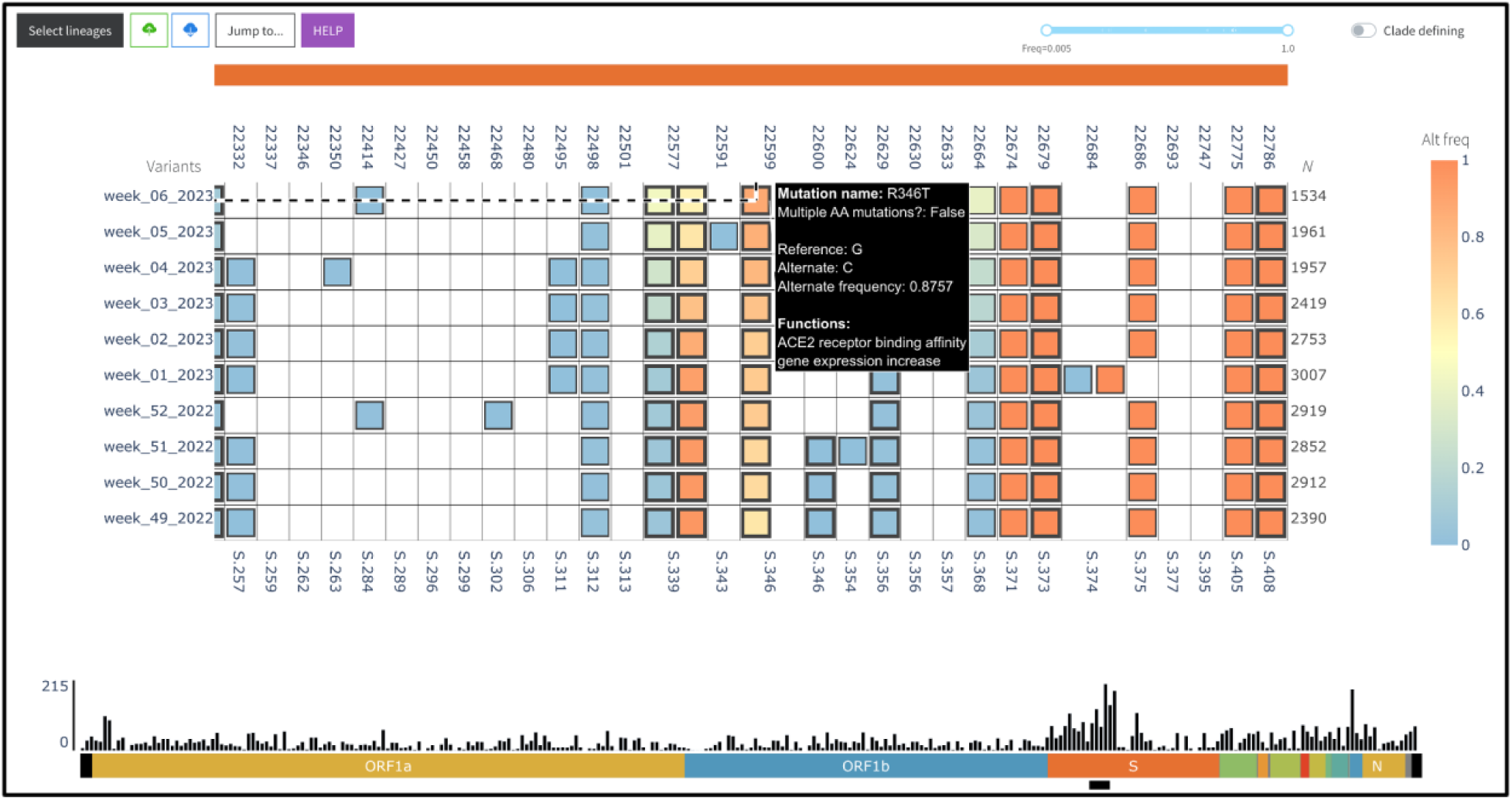
Longitudinal analysis of SARS-CoV-2 data from Canada (between Week 49 of 2022 and Week 06 of 2023) using VIRUS-MVP.

Applying this setup, we identified a notable increase in the S:R346T mutation in the spike protein, which appeared in Canada near the end of 2022. The S:R346T mutation enhances the virus’s binding affinity for ACE2 receptors on host cells, which can increase infection potential and impact virulence (42). This mutation is prevalent in the Omicron sub-lineage XBB.1.5, initially detected in Canada around October 2022. The swift rise in XBB.1.5 cases corresponded with the increased frequency of S:R346T, underscoring VIRUS-MVP’s ability to identify mutations of interest early in their emergence.

By monitoring the growth trajectory of this mutation and its eventual establishment within an epidemiologically significant lineage, we gained insights into its potential selective advantage and fitness implications and demonstrated how VIRUS-MVP could support proactive genomic surveillance. This capability is particularly valuable for public health stakeholders, as it allows for early detection of mutations that may be associated with increased transmission or disease severity, potentially shaping research and intervention priorities.

#### 3.2.2 Integrated wastewater and WGS analysis

Sequencing of viral genomes has been pivotal in COVID-19 surveillance, supporting pandemic response and enabling the study of viral evolution and prevalence. As part of a sustainable, long-term approach, a One-Health surveillance model is being implemented to integrate environmental surveillance with clinical monitoring systems (43, 44). Environmental surveillance, particularly wastewater sampling, allows the detection of viral variants and mutations of interest within the community, providing early signals of viral dissemination across broader populations. Using VIRUS-MVP, wastewater samples— sequenced independently with a dedicated workflow—can be analyzed for mutations and variant prevalence, then directly compared with clinical genome sequences within the same visualization. This comparison helps determine if detected mutations in wastewater samples also appear in clinical specimens, enhancing mutation tracking. To validate this capability, we analyzed simulated wastewater amplicon sequencing data (Sutcliffe et al., 2023) alongside real genome sequencing data from the Canadian VirusSeq Data Portal in the same analysis view (Figure 6). The use of simulated wastewater data allows us to know the ground truth.

**Figure 6.**
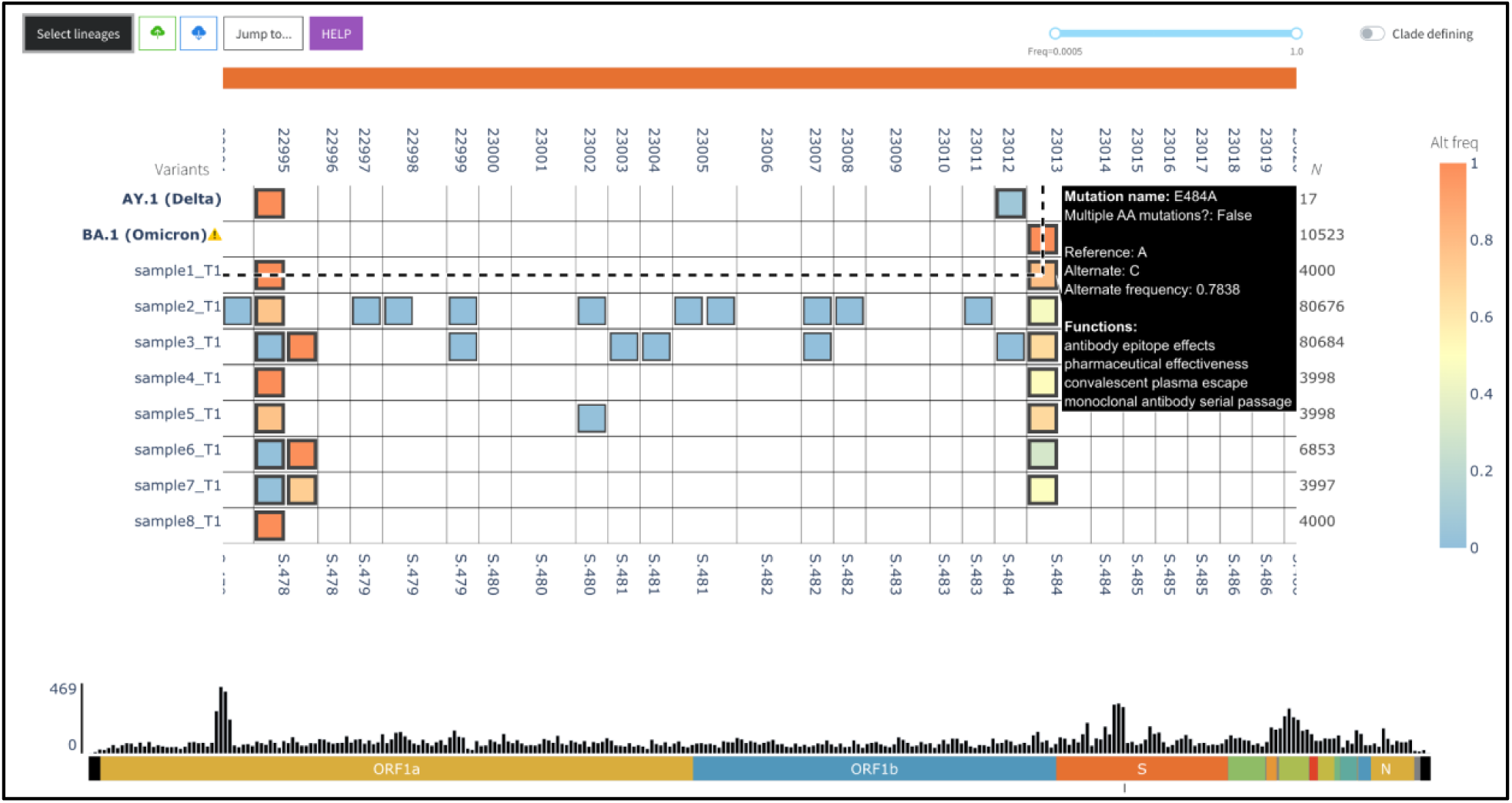
VIRUS-MVP adapted to visualize SARS-CoV-2 wastewater sequencing data and compared to the Canadian genome sequencing data from VirusSeq Data Portal in the same view

For example, in a simulated sample (sample1_T1) composed of a 75:25 lineage ratio of BA.1 (Omicron) to AY.1 (Delta), VIRUS-MVP identified the S:E484A mutation with a 78% prevalence. Notably, S:E484A is specific to BA.1 (Omicron) and absent in AY.1 (Delta), supporting the simulated lineage ratio. This approach demonstrates VIRUS-MVP’s capability for early detection of key mutations in wastewater samples, which can be validated in clinical samples to monitor ongoing viral evolution.

## 4. Discussion and future work

Molecular surveillance has been instrumental in tracking the evolution and spread of viruses, particularly during the COVID-19 pandemic. The unprecedented volume of genomic sequencing data available through global databases such as GISAID, Pathoplexus (45), the COVID-19 Data Portal at NCBI, the European Bioinformatics Institute (EBI), and national portals like the Canadian VirusSeq Data Portal(46) has enabled the development of numerous tools for analyzing SARS-CoV-2 lineages and variants. While impactful, these tools differ in their capabilities, emphasizing various aspects of genomic epidemiology such as mutation tracking, lineage assignment, and functional annotation.

VIRUS-MVP contributes to the shared aim of advancing genomic epidemiology by providing a flexible and portable genomics workflow that complements other tools (e.g. see Table 1) in the ecosystem which generally does not accept user data (10–12, 47). While platforms such as CovMT(10) or CoV-Spectrum(11) focus on real-time mutation tracking using predefined datasets to drive specific functionalities like variant proportion analysis, VIRUS-MVP offers unique value by enabling users to upload custom genomic datasets for real-time processing and visualization. This adaptability, combined with its pathogen-agnostic design, ensures that VIRUS-MVP can be tailored to diverse research needs and viral contexts. Rather than replacing existing viral surveillance dashboards, VIRUS-MVP aims to provide both data processing and visualization tools to the users for local data processing while also supporting the dashboard function on our website.

**Table 1.** Comparison of key features of mutation-centric analysis across different tools and platforms.

| Feature | CovRadar | Cov-MT | CoV-Spectrum | Outbreak.info | VIRUS-MVP |
| --- | --- | --- | --- | --- | --- |
| Mutation Focus | Spike gene mutations | All Mutations | All Mutations | All Mutations | All Mutations |
| Data Source | Global sample collections | Global sample collections | Global sample collections | Global sample collections | Customizable dataset (user-defined sources) |
| Visualization | Frequencies and spatiotemporal distributions | Evolution of mutational landscape | Proportion of different variants over time and space | Web-based visualization dashboard for exploring mutational content and lineage information | Interactive visualization of mutations, variants, and their functional impact |
| Functional Annotations | Not available | Summarizes mutations based on severity | Provides information on hospitalization , and mortality probabilities | Provides literature linked to lineages and location-based prevalence | Integrates curated functional annotations and supports overlaying custom annotations |
| Accessibility | Web-based platform | Web-based platform | Web-based platform | Web-based platform | Web-based and local deployment |
| Open Source | Yes (GPL-3.0 License) | Yes (MIT License) | Yes (GPL-3.0 License) | Yes (GPL-3.0 License) | Yes (MIT License) |
| Customization | Limited | Limited | Limited | Limited | Highly customizable (configurable workflows, annotation layers) |
| Integration | Stand-alone tool | Stand-alone tool | Stand-alone tool | - | Possibility to use it as a stand-alone or Integrated framework |

One of VIRUS-MVP’s key contributions is its robust functional annotation pipeline. It integrates curated insights into viral mutations, their potential phenotypic effects, and standardized external annotation overlays. This complements existing platforms such as Outbreak.info(47), which links mutations to literature and clinical outcomes, by offering an integrated, highly customizable annotation framework. Together, these tools enrich the landscape of genomic epidemiology, providing researchers with a comprehensive toolkit to analyze and interpret viral genomes.

As part of Canada’s national viral surveillance ecosystem, a production instance of VIRUS-MVP has been deployed for SARS-CoV-2 surveillance. This instance leverages WGS data and associated metadata from the Canadian VirusSeq Data Portal, enabling weekly updates with newly submitted sequences from public health laboratories across the country. For mpox, a production instance is also available at virusmvp.org/mpox, utilizing data from PathoPlexus. These deployments exemplify VIRUS-MVP’s adaptability to diverse pathogens and research needs.

Looking ahead, VIRUS-MVP aims to expand its capabilities and application scope. Ongoing development includes integrating additional viral pathogens such as RSV, Influenza, and HIV. Addressing challenges such as data curation bottlenecks will involve leveraging large language models to automate annotation and metadata extraction. Furthermore, we encourage community contributions via the platform’s GitHub repository (https://github.com/cidgoh/VIRUS-MVP), where researchers can propose new workflows, modules, and features.

VIRUS-MVP has evolved from a COVID-19-specific tool to a pathogen-agnostic platform with wide-reaching potential in infectious disease research and surveillance. Its unique combination of customizable workflows, real-time visualization, and integrated functional annotations positions it as a critical resource for public health practitioners, epidemiologists, and researchers worldwide. By continuing to refine its capabilities and expand its viral repertoire, VIRUS-MVP aims to play a pivotal role in addressing both current and emerging viral threats.

## 5. Availability and Deployment

We have deployed VIRUS-MVP on a cloud service using the docker-compose setup found on the application GitHub repository. The VIRUS-MVP docker image is built on top of the latest Python base image, and the resulting container is served through gunicorn (48). To establish a secure SSL connection, we implemented a Nginx reverse proxy setup (49) external to the VIRUS-MVP GitHub repository.

For users seeking to deploy an application instance with upload functionality, the genomics workflow built within the deployed container also utilizes a Docker containerization strategy by default. This strategy was implemented due to the relatively slower performance of a Conda containerization strategy, and difficulties with installing Singularity inside an existing Docker container. To avoid a Docker-in-Docker scenario leading to container escape, the docker-compose.yml file bind mounts the host docker socket. This Docker-based architecture makes VIRUS-MVP highly portable across various cloud environments. With containerized components, users can deploy VIRUS-MVP on any cloud provider that supports Docker, enabling flexible scaling and straightforward replication in new environments.

1. VIRUS-MVP instance dedicated to SARS-CoV-2 data - https://virusmvp.org/covid-mvp
2. VIRUS-MVP visualization source code, manual, and deployment - https://github.com/cidgoh/VIRUS-MVP
3. VIRUS-MVP Genomics workflow source code - https://github.com/cidgoh/nf-ncov-voc
4. Functional Database Pokay - https://github.com/nodrogluap/pokay
5. Mutation Functional Annotation Contextual Data Specifications package https://github.com/cidgoh/pathogen-mutation-functionalannotation-package
6. SARS-CoV-2 surveillance indicators for public health - https://github.com/cidgoh/nf-ncov-voc/tree/master/assets/ncov_surveillanceIndicators
7. Surveillance report (TSV format description) - https://github.com/cidgoh/nf-ncov-voc/blob/master/docs/surveillance_report_description.md
8. Surveillance report (PDF format example) - https://github.com/cidgoh/nf-ncov-voc/blob/master/docs/Omicron_surveillance_report.pdf
9. All sequence data used in developing, testing, and generating the figure is available at https://virusseq-dataportal.ca

## Declarations

### Ethics approval and consent to participate

Not Applicable

## Acknowledgments

The results here are in whole or in part based upon data hosted at the Canadian VirusSeq Data Portal: https://virusseq-dataportal.ca/. We acknowledge the Canadian Public Health Laboratory Network (CPHLN), Genome Canada, and the CanCOGeN VirusSeq Consortium for their valuable contributions to the Portal; see the supplementary file for detailed information. We also thank the Computational Analysis Modelling and Evolutionary Outcomes (CAMEO) initiative of the Coronavirus Variants Rapid Response Network (CoVaRR-Net) for their regular feedback. Additionally, we are grateful to the CPHLN Genomics Work Group, particularly Inès Levade and Keith Mackenzie, for their extensive feedback on this work. We are also grateful to the PHA4GE Data Structures Working Group for their insightful feedback, which improved our data organization approach.

## Authors’ contributions

**Conceptualization:** W.W.L.H., M.Z.A., I.S.G. **Software development:** M.Z.A., I.S.G., M.I., A.S. **Data curation:** P.M.K.G., M.I., A.S., K.V., F.S.L.B. **Data standardization and harmonization:** M.I., M.Z.A., P.M.K.G., E.J.G., D.D., W.W.L.H. **Deployment:** J.D., I.S.G., M.Z.A. **Writing - original draft:** M.Z.A. **Writing - review and editing:** M.Z.A., I.S.G., M.I., A.S., E.J.G., D.D., J.D., K.V., G.V.D., F.S.L.B., P.M.K.G., W.W.L.H. **Project administration:** M.Z.A. **Supervision:** G.V.D., F.S.L.B., P.M.K.G., W.W.L.H. **Funding:** W.W.L.H.

